# Does genetic differentiation underlie behavioral divergence in response to migration barriers in sticklebacks? A common garden experiment

**DOI:** 10.1101/2021.08.25.457647

**Authors:** A. Ramesh, M. M. Domingues, E.J. Stamhuis, A.G.G. Groothuis, F.J. Weissing, M. Nicolaus

## Abstract

Water management measures in the 1970s in the Netherlands have produced a large number of ‘resident’ populations of three-spined sticklebacks that are no longer able to migrate to the sea. This may be viewed as a replicated field experiment, allowing us to study how the resident populations are coping with human-induced barriers to migration. We have previously shown that residents are smaller, bolder, more exploratory, more active, more aggressive, exhibited lower shoaling and lower migratory tendencies compared to their ancestral ‘migrant’ counterparts. However, it is not clear if these differences in wild-caught residents and migrants reflect genetic differentiation, rather than different developmental conditions. To investigate this, we raised offspring of four crosses (migrant ♂ x migrant ♀, resident ♂ x resident ♀, migrant ♂ x resident ♀, resident ♂ x migrant ♀) under similar controlled conditions and tested for differences in morphology and behavior as adults. We found that lab-raised resident sticklebacks exhibited lower shoaling and migratory tendencies as compared to lab-raised migrants, retaining the differences in their wild-caught parents. This indicates genetic differentiation of these traits. For all other traits, the lab-raised sticklebacks of the various crosses did not differ significantly, suggesting that the earlier-found contrast between wild-caught fish reflect differences in their environment. Our study shows that barriers to migration can lead to rapid differentiation in behavioral tendencies over contemporary timescales (∼50 generations), and that part of these differences reflects genetic differentiation.

**Significance statement:** Many organisms face changes to their habitats due to human activities. Much research is therefore dedicated to the question whether and how organisms are able to adapt to novel conditions. We address this question in three-spined sticklebacks, where water management measures cut off some populations, prohibiting their seasonal migration to the North Sea. In a previous study, we showed that wild-caught ‘resident’ fish exhibited markedly different behavior than migrants. To disentangle whether these differences reflect genetic differentiation or differences in the conditions under which the wild-caught fish grew up, we conducted crosses, raising the F1 offspring under identical conditions. As their wild-caught parents, the F1 of resident x resident crosses exhibited lower migratory and shoaling tendencies than the F1 of migrant x migrant crosses, while the F1 of hybrid crosses were intermediate. This suggests that ∼50 years of isolation are sufficient to induce behaviorally relevant genetic differentiation.

## Introduction

Habitat fragmentation resulting from human activities is considered to be a major threat for many animal populations (Foley et al. 2005; Fischer and Lindenmayer 2007). Habitat fragmentation is characterized by a reduction in habitat size, habitat loss, and loss of habitat connectivity (Fahrig 2003). This poses a threat to animal populations, especially for migratory species which rely on connectivity between functional habitats for reproduction and survival (Legrand et al. 2017). Migratory species would thus need to respond via adaptive changes in life history and behavior to thrive in disconnected patches (Bohlin et al. 2001; Kraabøl et al. 2009; Junge et al. 2014). Therefore, understanding the underlying mechanisms of these responses is crucial as they directly affect the future adaptive potential and evolutionary trajectories of populations (Kawecki and Ebert 2004; Wang and Bradburd 2014) as well as conservation measures (Stockwell et al. 2003).

Individuals need to maintain a match between their phenotypes and the environment to enhance their local performance, thereby allowing populations to subsist or grow in an altered environment. Depending on the underlying mechanism involved, such adaptive responses may occur more or less rapidly and may influence population genetic structure (Hedrick et al. 1976; Hedrick 2006; Nicolaus and Edelaar 2018). For example, phenotypic adjustment may result from natural selection favoring some phenotypes over others, potentially leading to population genetic differentiation across multiple generations when phenotypic variation has a genetic basis (Kawecki and Ebert 2004). Non-exclusively, individuals may match their phenotype to local conditions through plasticity, be it reversible plasticity (or phenotypic flexibility *sensu* Piersma and Drent 2003), developmental plasticity, or transgenerational plasticity (through parental and epigenetic effects). Plasticity, defined as the ability of a genotype to exhibit different phenotypes in response to the environment (Via et al. 1995; Pigliucci 2005), can thus provide a rapid mechanism to respond to environmental changes (Ghalambor et al. 2007). Importantly, selection may favor genotypes with varying levels of plasticity (Nussey et al. 2007, Scheiner 1993), implying that mentioned mechanisms are intertwined (Edelaar et al. 2017) and that observed population divergence could reflect genetic differentiation and/or differences in the environments under which individuals grow up. In migratory species, migrants would have to exhibit phenotypic plasticity or bet-hedging strategies, as they are exposed to different environmental conditions (Botero et al. 2015). In the case where migrants are no longer able to migrate (forced ‘residents’), we expect selection to act on either the traits themselves or on the degree of plasticity.

In this study, we focus on behavior as it is the primary way through which animals interact with their environment and respond to changes (Wong and Candolin 2015). Behavior is often considered highly flexible and hence less prone to genetic divergence in response to environmental changes. However, plastic responses could evolve rapidly through genetic divergence compared to fixed traits (Van Gestel and Weissing 2018). In addition, ‘animal personality’ research points that behaviors are highly structured and form correlations over time (consistency) and over contexts (syndromes) (Réale et al. 2007; Stamps and Groothuis 2010; Wolf and Weissing 2012). Furthermore, individual differences within populations are often repeatable (Réale et al. 2007) and to some extent, heritable (Dochtermann et al. 2014). As a consequence, personality variation may retard or accelerate rates of microevolution and population divergences (Wagner and Altenberg 1996 ; Wolf and Weissing 2012; Dochtermann and Dingemanse 2013; Van Gestel and Weissing 2018). Here we aim to study whether genetic differentiation underlies the rapid behavioral differentiation following habitat fragmentation. We capitalize on an unintended field experiment in the north of The Netherlands, where the construction of pumping stations in the 1970s has led to the forced residency of replicate populations of anadromous three-spined sticklebacks (*Gasterosteus aculeatus*). A previous study in this system has revealed extensive phenotypic differentiation (morphology and behavior) between the ancestral ‘migrant’ and its derived ‘resident’ populations (Ramesh et al. 2021). Compared to migrants, wild-caught residents are smaller, more active and aggressive, more exploratory, bolder, and showed reduced shoaling and migratory tendencies (Ramesh et al. 2021). These differences parallel the behavioral divergence reported between freshwater and marine populations of sticklebacks over ∼12,000 years (Di-Poi et al. 2014). However, it remains to be determined if similar behaviorally relevant genetic differentiation has evolved in our system over much shorter time scales (∼50 years). This knowledge is important because conservation efforts are underway to reconnect the waterways and therefore, we need to better understand the current state of fish populations in order to predict the eco-evolutionary consequences of barrier removal.

We conducted a common garden experiment to test whether genetic differentiation underlies the observed divergence in morphology and behavior. We raised F1 juveniles from four types of crosses (migrant parents (MM), resident parents (RR), hybrids with a migrant mother (RM) and hybrids with a resident mother (MR); Fig 1a) under similar laboratory conditions and quantified variation in activity, exploration, shoaling, boldness, and migratory tendencies among these crosses. We expect that 1) if the behavioral differentiation is genetic, individuals of MM crosses will differ significantly from RR crosses (similar to their wild-caught parents), 2) if the behavioral differences between wild-caught residents and migrants are induced by differences in their environments, there will be no differences between the ‘common garden’ crosses; and 3) if parental effects are involved, we will see asymmetric changes in the reciprocal hybrid crosses (Fig. 1b). Specifically, if behavioral variation is strongly influenced by maternal effects, the hybrids resulting from the MR cross will have a similar score as the RR cross and the hybrids resulting from the RM cross will have a similar score as the MM cross (Fig. 1b). A similar trend can be expected in the case of paternal effects, but we eliminated that possibility to a large extent by raising juveniles without paternal care (Giesing et al. 2011; McGhee and Bell 2014; Heckwolf et al. 2018).

**Fig. 1 a.**
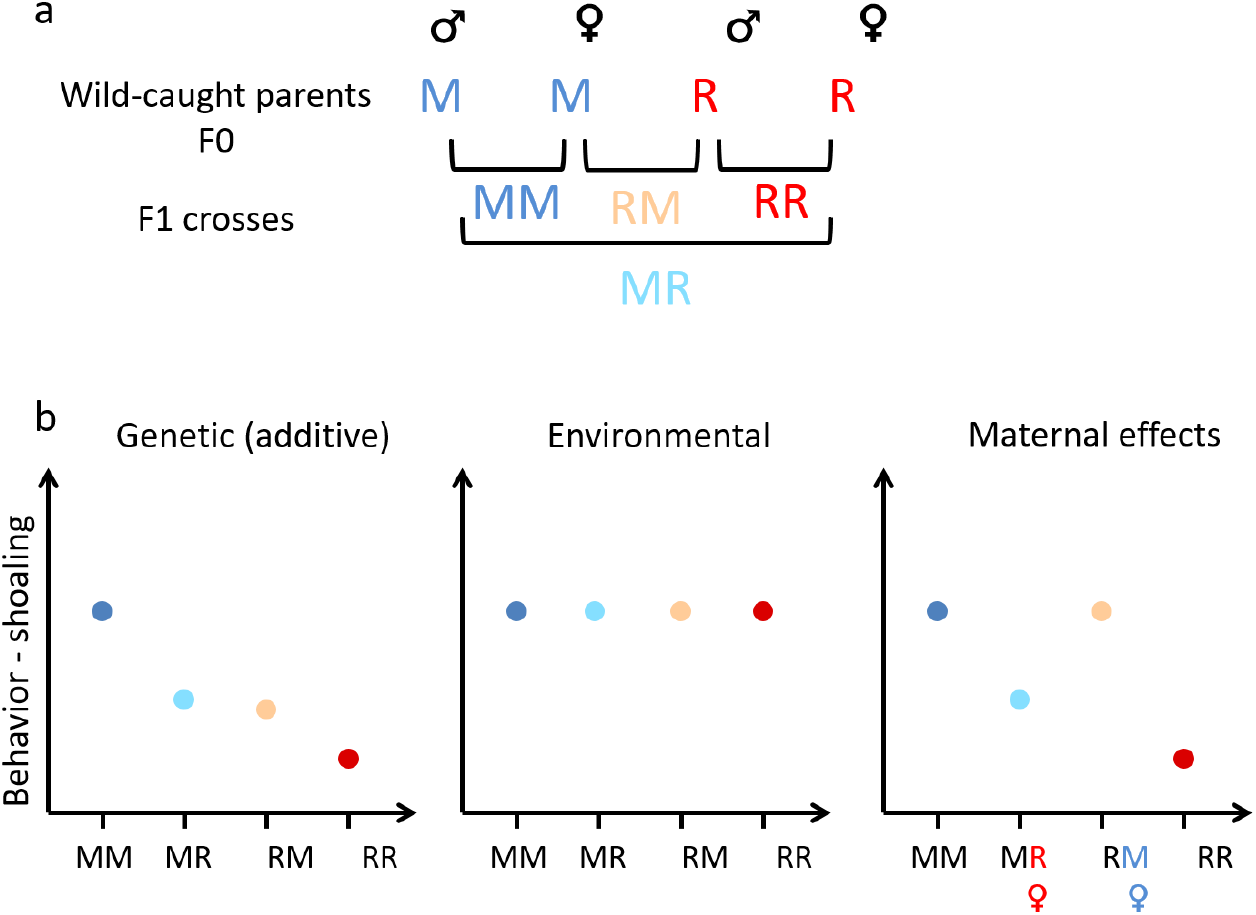
Schematic of breeding design. We obtained four F1 crosses – migrant male × migrant female (MM), resident male × resident female (RR), migrant male × resident female (MR) and resident male × migrant female (RM); **b** Expectations of mean behavioral scores (for example, shoaling) if the underlying basis for behavioral differentiation in wild-caught parents is due to genetic differentiation, environmental experiences during development or through maternal effects. (Letters of migrant and resident female in the maternal effects prediction plot are colored according to the origin for ease of interpretation of patterns in hybrids, when they are under the control of maternal effects. The expected mean value of hybrids, would correspond to the migrant or resident status of the female.)

## Methods

### Study populations

The waterways in the Netherlands consist of rivers and canals that are open to the sea and of land-locked smaller ditches (<1 m deep) located in side polders. We caught incoming migrants at two sea locks (“TER” (*53^0^18’7.24’’, 7^0^2’17.11’’*) and “NSTZ” (*53^0^13’54.49’’, 7^0^12’30.99’’*)) whereas residents were caught in two land-locked polders (“LL-A” (*53^0^17’56.14’’, 7^0^2’1.28’’*) and “LL-B” (*53^0^17’16.52’’, 7^0^2’26.46’’*)) (Ramesh et al. 2021). Sticklebacks were caught over a period of four weeks between March and April in 2019. All individuals were transported to the laboratory within two hours of capture in aerated bags (5-6 fish / 3L bag). They were housed outdoors separated by their origin in groups of five fish in 50 liter aerated tanks filled with freshwater, exposed to the natural day-light cycles and temperatures. They were fed brine *ad libitum* with brine shrimps and blood worms (3F Frozen Fish Food bv.). Males were separated once they reached breeding colors, and females were checked daily for signs of gravidity.

### Lab-bred F1 juveniles

Lab-bred F1 juveniles of resident, migrant, and hybrid sticklebacks arose from a partial factorial breeding design (Fig. 1a) using three resident males, three resident females, three migrant males, and three migrant females (six migrants from “NSTZ”, five residents from “LL-A” and one resident female from “LL-B”). Each family consisted of all combinations of crosses between a male and female migrant and male and female resident, leading to F1 offspring of different crosses: pure migrant (MM) or resident (RR) and hybrids with migrant father and resident mother (MR) and vice versa (RM). From the offspring pool, a total of 40 fish were used per cross for the experiment, with each cross containing at least five fish from each family.

For obtaining F1 juveniles, we followed a split-clutch in-vitro fertilization protocol, where eggs of ripe females were stripped, then weighed and split into two halves for artificial insemination with sperm extracted from freshly euthanized migrant and resident fathers respectively (Barber and Arnott 2000). All offspring were raised without paternal care to prevent undesired long-lasting effects of father on offspring behavior (McGhee and Bell 2014). The larvae hatched five to seven days after fertilization and started maintaining buoyancy and independent feeding one week after hatching. The fish larvae were fed a mixture of frozen cyclops, freshly hatched Artemia nauplii, and zebrafish diet (GEMMA Micro 75, Skretting, Tooele, Utah) daily. The densities never exceeded 40 fish larvae in 5 liter “home-tanks” (30 × 16 × 18 cm (L × W × H)). Once fish reached ∼2 cm, they were isolated, assigning ten random individuals from the same family into separate home tanks. After this, the individuals were fed *ad libitum* with brine shrimps and blood worms (3F Frozen Fish Food bv.), and tanks were connected to the same water system at 16° C. The photoperiod was set at 16:8 (L:D), mimicking summer conditions during juvenile growth. When the fish reached a length of ∼4 cm, they received a unique identification (see below). We induced autumn conditions when the fish were ∼12-13 months old, characterized by 12:12 (L:D) photoperiod and temperatures being lowered to 13° C – 14° C. All fish were in non-breeding conditions and kept in autumn conditions during the period of experimentation. Experimentation started when fish were ∼15-16 months old.

### Individual identification

When the juveniles reached 4cm length (∼12 months), we used clipped spines or injection of an 8 mm Passive Integrated Transponder (PIT tag; Trovan, Ltd., Santa Barbara, California) for unique individual identification, We used PIT tag injection only for half of the tested fish (20 fish X 4 crosses = 80 fish), while the rest were tagged using a combination of dorsal and pelvic spine clipping (20 fish X 4 crosses = 80 fish). This was because PIT tag retention was low in these fish (∼15% loss in the first week after tagging) and we did not retag the fish to prevent excess handling. PIT tags were injected in the abdominal cavity and under anesthesia following the standard protocol (following Cousin et al. 2012). During tagging/clipping, we also measured weight and standard length (the length from the tip of the snout to the base of the tail) as a proxy for size. Lateral plates were not very clearly visible in juvenile fish and hence were not measured. After individual tagging, we mixed juveniles from different families to be housed together in groups of ten in their home tanks while keeping them together with the same cross (MM, RR, MR or RM).

### Large-scale movement tendencies in mesocosm (migratory tendencies)

For the subset of PIT tagged fish, movement assays were performed in semi-natural mesocosms before subjecting them to the lab-based tests. The mesocosm system consisted of five connected outdoor ponds of diameter 1.6m connected by four pipes of length ∼1.5 m and diameter 11cm, filled with water from a nearby freshwater ditch, with a linear flow in the system of connected ponds similar to those typically experienced in the canals and ditches (flow speed < 0.7 cm/s) (Fig 2a). This was done to create a cue for migration-like movement. All connecting tubes were fitted with circular PIT antennas around the entrance and exit of each pond to record fish movement between ponds. The sticklebacks were tested in pond experiments after at least one week of recovery from tagging. A group of ten fish of one cross (MM, RR, MR or RM) was introduced in the first pond and acclimatized for 5 hours in the first morning, after which the connection to the rest of the ponds was opened. We then recorded the movement of fish as the number of crossings between ponds for the next 16 h (∼4 p.m.-8 a.m.). We attempted to have 20 tagged fish/cross and tested them in groups of 10 each, making it 2 groups/cross. However, due to tag loss, we ended up with <20 fish/cross. Instead of changing group size, which could have an effect on behavior, we decided to spread the final number of tagged fish between two groups and supplement the remaining with untagged fish from the same cross to make up to 10. In total, 2 groups, each from a randomly chosen cross, were tested, making a total of eight groups with 56 fish (N_MM_ = 12, N_MR_ = 17, N_RM_ = 15, N_RR_ = 12). In groups with less than ten tagged individuals, untagged fish from the same cross were added to maintain constant group size.

**Fig. 2.**
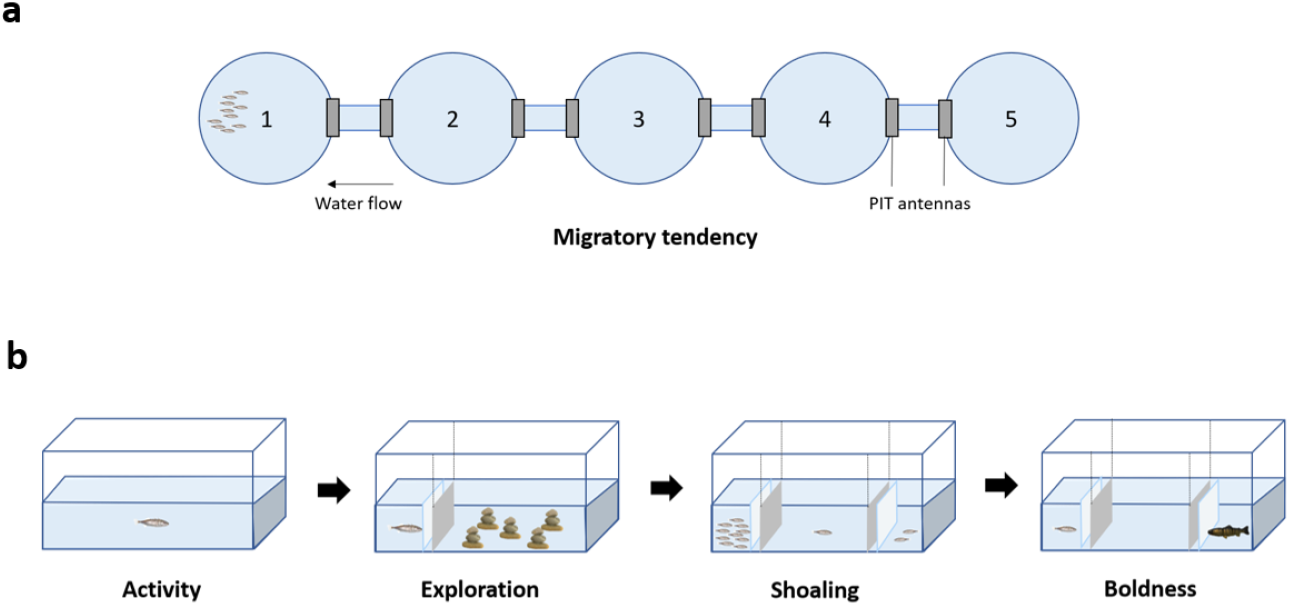
Schematic of behavioral assays. **a** Migration tendencies were tested in a linear setup of five connected pond mesocosm with groups of 10 fish. There is water flow (rate < 0.7 cm/s). PIT antennas are present at both ends of the corridors connecting the ponds **b** Lab assays were performed in the following order: Activity (Day 1), Exploration (Day 1), Shoaling (Day 2), Boldness (Day 3)

### Lab behavior assays

Three days before testing, fish were selected randomly and acclimatized in visually separated and isolated tanks, same as their home tanks, at an ambient temperature of 19°C. We attempted to test 40 fish/cross, but some fish were lost due to mortality. Hence in total, 154 fish (N_MM_ = 40, N_MR_ = 39, N_RM_ = 35, N_RR_ = 40) were randomly selected for testing and split into four batches. One round of testing consisted of four batches of ∼10 fish of each cross and lasted one week where we assayed activity, exploration, shoaling, and boldness in that order (Fig 2b). Overall, roughly 40 fish were tested each week. The interval between the first and the second round of testing of each individual was thus at least four weeks. Fish were returned to their home tanks between the testing rounds.

The sample sizes for the second round was lower (N = 151) due to mortality between the two rounds (N_MM_ = 39, N_MR_ = 38, N_RM_ = 34, N_RR_ = 40). All lab-assays were filmed from the top using a Raspberry Pi camera (Raspberry Pi NoIR Camera Board V2 – 8MP, Raspberry Pi Foundation, UK) in tanks placed in illuminated wooden boxes to prevent external disturbance. Behavioral assays were conducted in fixed order as below, and videos were analyzed using EthovisionXT (Noldus Information Technology bv.). In all tests, observers were blind with respect to the cross to which the test fish belonged and further bias was reduced by analyzing the videos using automated video tracking techniques.

#### Activity (day 1)

Activity of the fish was measured as the total distance the fish swam in a tank identical to its home tank during a total of 20 min (with 5 min for acclimatization).

#### Exploration (day 1)

Just after activity was recorded, the fish was isolated to one corner of the tank using a sheet partition, and the setup in the tank was changed. Five stone pillars extending above the water’s surface were added in a specific position, forcing the fish to move around them. After 5 min, the sheet was removed remotely without opening the box, and the fish was recorded in this novel arena for 20 min. The total distance travelled by the fish in this novel environment was used as a proxy for exploratory tendency of fish as it highly correlates with space use (Ramesh et al. 2021).

#### Shoaling (day 2)

For the shoaling assay, a larger tank (60 × 30 × 30 cm) was filled with water up to 10 cm height. The tank was divided into three compartments: the central testing arena where the focal fish was released and two end compartments containing the stimulus shoal (n=10 unfamiliar conspecifics of mixed crosses), and the distractor fish (n=2 unfamiliar conspecifics) (adapted from Wark et al. 2011). The position of the distractor and shoal fish compartments was switched to prevent biases and replaced with new distractor and shoal fish every seven tests. At the start of the test, the focal fish was allowed to acclimatize for 5 min in the central arena without viewing the end compartments which were covered with opaque barriers. Then the opaque barriers were lifted remotely from outside the box, and the response of the focal fish was recorded for the next 20 min. The water was refreshed after testing seven fish in the arena. In total, we had four groups of shoal fish and five pairs of distractor fish, which were randomly used to avoid biases. The proportion of time the focal fish spent within one-fish distance (6 cm) from the side containing the stimulus shoal was used as a proxy for shoaling.

#### Boldness (day 3)

In the boldness tests, we measured the responses of the focal fish toward visual cue of an European perch (*Perca fluviatilis*) (model with soft body, Kozak and Boughman 2012) and olfactory predation cues (50 ml of water from freshly dead sticklebacks mixed with 50 ml of water containing live perch scent, Sanogo et al. 2011). The focal fish was moved from its home-tank into a bigger, novel tank (60 × 30 × 30 cm) with three compartments filled with 10 cm of water. The predator model was randomly presented in one of the end compartments, while the focal fish was acclimatized in the other end compartment (Kozak and Boughman 2012). After 5 min of acclimatization, the fish was released remotely into the arena with view of the predator model, and the assay lasted for 20 min. We changed the side of predator compartment systematically in order to avoid biases. Further, the water was refreshed and new predatory olfactory cues were added after testing seven fish in the arena. The proportion of time the focal fish spent within one-fish distance (6 cm) from the predator compartment was taken as a proxy for boldness.

### Statistical analyses

Variation in size and behaviors (activity, exploration, shoaling, and boldness) was analyzed using Linear mixed models (LMM) in which repeat (first vs. second round) and cross identity (MM, MR, RM or RR) were included as fixed factors. We also included the interactive effects (cross x round) to test for cross-specific habituation effects. Individual identity (Fish ID), mother identity (Mother ID) and father identity (Father ID) were included as random effects. For shoaling behavior, we added identity of the test shoal (Shoal ID) as an additional random effect. For migratory tendencies, only one round of tests was performed and we fitted a Poisson generalized linear mixed model with log-link function (GLMM), with number of pond crosses as the response variable and cross identity as fixed factor. As random effects, we included mother identity (Mother ID) and father identity (Father ID) and further, to prevent overestimation of predictive power caused due to overdispersion we added observation level random effects (OLRE) (Harrison 2014).

All LMMs/GLMMs were constructed in R v. 3.6.1, R Core Team (2019) using the ‘lmer’ function of the ‘lme4’. Package version 1.1-27.1 (Bates et al. 2015). The statistical significance of fixed effects was assessed based on the 95% confidence interval (CI): an effect was considered significant when its 95% CI did not include zero. In addition, Tukey’s HSD post-hoc test was performed using the functions ‘emmeans’ and ‘pairs’ to give pairwise comparisons using the package ‘emmeans’, package version 1.6.1 (Lenth 2020). LMMs were used to decompose the phenotypic variance of behaviors into between-individual (V_Fish ID_), between-mother (V_Mother ID_), between-father (V_Father ID_) and within-individual (V_Residual_) variances that we subsequently used to calculate repeatabilities, i.e the proportion of total phenotypic variation (V_p_) attributable to differences between individuals (R_fish ID_), between mothers (R_Mother ID_) and between father (R_Father ID_):

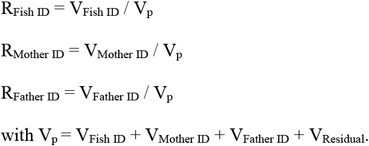

Raw (without fixed effects), adjusted repeatabilities (after accounting for fixed effects, cross x round), and their confidence intervals were calculated using ‘rpt’ function with 1000 bootstraps in ‘rptR’, package version 0.9.22 (Stoffel et al. 2017).

## Results

Our prime goal was to test if RR and MM crosses that were raised under similar conditions, exhibited similar behavioral differences as observed in their wild-caught population of origin and if these differences were consistent over time. We found that RR crosses were consistently less active than MM crosses in the two rounds (Fig. 3a; Table 1; overall effect of crosses on activity: χ^2^=17.35, df=3, p<0.01, Supp. Table 1). We further found that shoaling and migratory tendencies varied significantly and consistently between RR and MM crosses in the same direction, with RR crosses exhibiting lower shoaling and migratory tendencies than MM crosses (Fig. 3c, 2e; Table 1; overall effect of crosses on shoaling: χ^2^=17.91, df=3, p<0.01, on migratory tendency: χ^2^=14.37, df=3, p<0.01). MM but not RR crosses shoaled more than expected by chance (score of >0.5) (Table 1). RR and MM crosses did not differ consistently in levels of exploration and boldness (Fig. 3b, d; Table 1, Supp. Table 1). For boldness, RR cross differed from MM cross but only in round 2 (Fig. 3d, significant effects of round and round x cross RR, Table 1, Supp. Table 1), implying that the observed difference was not consistent over time (Fig. 3d). Crosses did not differ in body size (Fig. 3f, Table 1, Supp. Table 1).

**Table 1.** Effect of type of cross (migrant MM, resident RR, hybrid RM and MR) on behavior and morphology of common garden raised three-spined sticklebacks. For lab-based behaviors, the additive and/or interactive effects of rounds are included. Summaries of linear mixed models on traits are presented with estimates of fixed effects (β), with their 95% confidence intervals (CI) and variance due to random effects with corresponding standard deviation. Significant fixed effect compared to the reference factor are denoted in bold. The corresponding significant pair-wise comparisons (Tukey’s HSD post-hoc tests) are given in Fig. 2. Sample size (N) represents number of observations

|  | Activity<br>N = 305 | Exploration<br>N = 304 | Shoaling<br>N = 302 | Boldness<br>N = 303 | Migratory tendency<br>N = 52 | Size<br>N = 154 |
| --- | --- | --- | --- | --- | --- | --- |
| *Fixed effects* | $\beta$ (95% CI) | $\beta$ (95% CI) | $\beta$ (95% CI) | $\beta$ (95% CI) | $\beta$ (95% CI) | $\beta$ (95% CI) |
| **Intercept (Round First, Cross MM)** | **49.56 (44.15, 56.64)** | **9.11 (6.47, 11.76)** | **0.63 (0.57, 0.70)** | **0.55 (0.47, 0.63)** | **2.86 (2.07, 3.62)** | **47.01 (44.98, 49.04)** |
| **Cross (MR)** | -4.51 (-16.10, 7.03) | -1.79 (-5.53, 1.92) | **-0.10 (-0.18, 0.03)** | -0.03 (-0.12, 0.07) | -0.51 (-1.63, 0.54) | -0.79 (-2.86, 1.21) |
| **Cross (RM)** | **-8.28 (-15.07, -0.29)** | -1.86 (-4.63, 1.14) | **-0.09 (-0.17, -0.01)** | -0.03 (-0.15, 0.08) | **-0.98 (-1.92, -0.04)** | 0.54 (-2.52, 3.28) |
| **Cross (RR)** | -5.54 (-17.05, 5.98) | -0.55 (-4.25, 3.14) | **-0.10 (-0.17, -0.02)** | -0.02 (-0.13, 0.09) | **-2.03 (-3.20, -0.86)** | -1.35 (-4.18, 1.51) |
| **Round (Second, Cross MM)** | **-8.66 (-13.93, -3.42)** | 0.91 (-1.21, 3.02) | 0.03 (-0.03, 0.10) | **0.11 (0.03, 0.19)** | - | - |
| **Cross (MR) X Round (Second)** | -4.91 (-12.04, 2.26) | -0.93 (-3.80, 1.95) | 0.004 (-0.09, 0.10) | 0.00 (-0.11, 0.11) | - | - |
| **Cross (RM) X Round (Second)** | **-11.62 (-19.13, -4.09)** | -2.31 (-5.33, 0.71) | -0.78 (-0.18, 0.03) | -0.08 (-0.20, 0.03) | - | - |
| **Cross (RR) X Round (Second)** | -5.35 (-12.48, 1.81) | -0.02 (-2.88, 2.87) | -0.02 (-0.12, 0.07) | **-0.12 (-0.23, -0.01)** | - | - |
| *Random effects* | Estimate (std.dev) | Estimate (std.dev) | Estimate (std.dev) | Estimate (std.dev) | Estimate (std.dev) | Estimate (std.dev) |
| **Fish ID** | 80.05 (8.97) | 13.10 (3.62) | 0.007 (0.09) | 0.01 (0.12) | - | - |
| **Father ID** | 0.00 (0.00) | 0.00 (0.00) | 0.001 (0.03) | 0.00 (0.05) | 0.00 (0.00) | 2.21 (1.49) |
| **Mother ID** | 41.69 (6.46) | 3.39 (1.84) | 0.00 (0.00) | 0.00 (0.02) | 0.08 (0.28) | 0.59 (0.77) |
| **Shoal ID** | - | - | 0.00 (0.03) | - | - | - |
| **OLRE** | - | - | - | - | 1.20 (1.10) | - |
| **Residual** | 111.7 (10.56) | 18.04 (4.25) | 0.02 (0.13) | 0.03 (0.16) | - | 11.21 (3.35) |

**Fig. 3.**
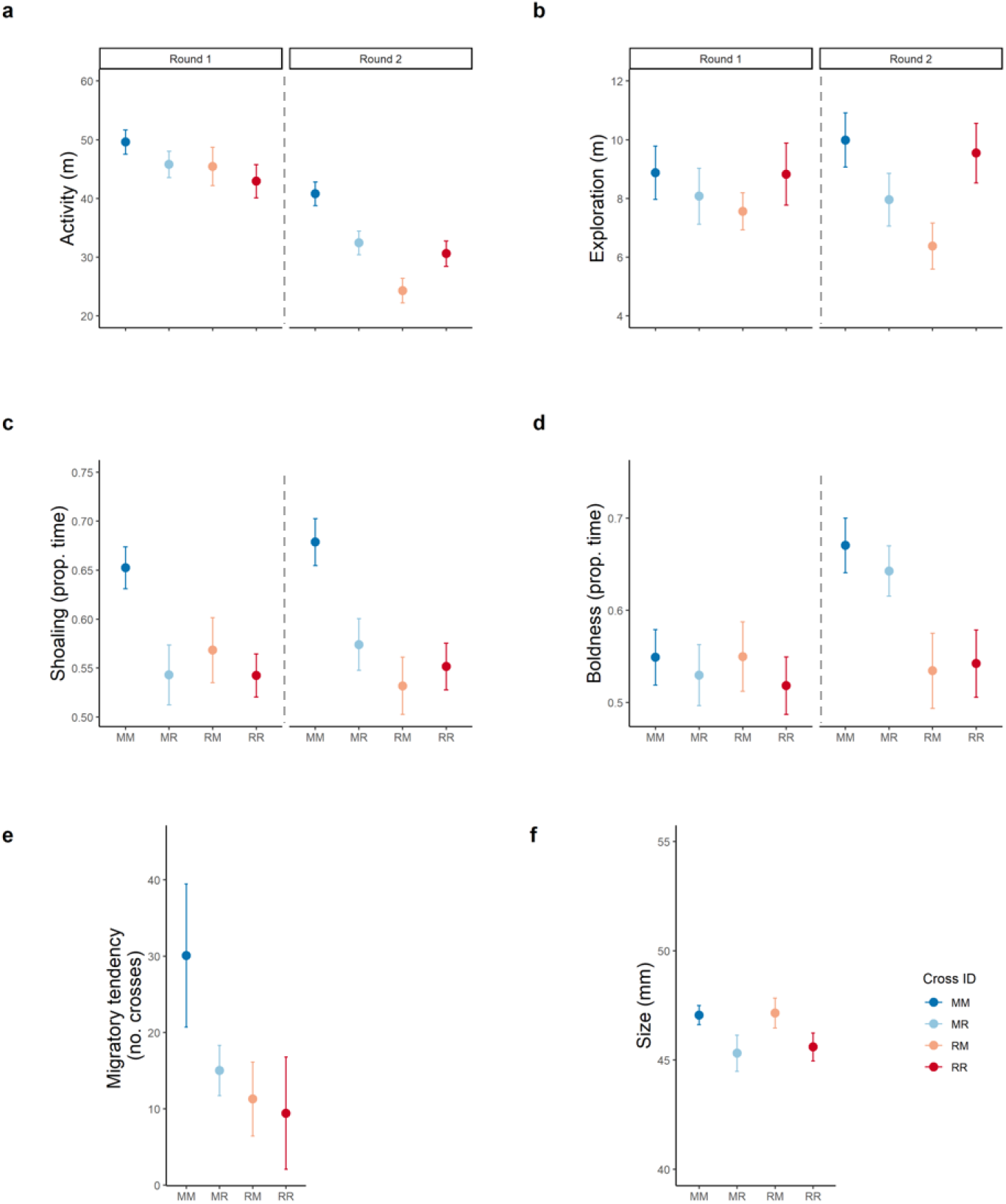
Mean scores and standard errors for behaviors and size of F1 fish of different crosses. **a** “Activity” – total distance travelled in meters (m); **b** “Exploration” – total distance travelled in a novel arena in m; **c** “Shoaling” – Proportion of time spent near shoal compartment; **d** “Boldness” – Proportion of time spent near predator. Means along with standard error are represented in the plots. For lab-based behaviors, the mean behavioral scores for the two repeats are represented separately. (Sample sizes round 1: N_MM_ = 40, N_MR_ = 39, N_RM_ = 35, N_RR_ = 40; round 2: N_MM_ = 39, N_MR_ = 38, N_RM_ = 34, N_RR_ = 40); **e** “Migratory tendency” – total number of pond crosses (N_MM_ = 12, N_MR_ = 17, N_RM_ = 15, N_RR_ = 12); **f** “Size” – Standard length in mm (N_MM_ = 40, N_MR_ = 39, N_RM_ = 35, N_RR_ = 40).

We did not find evidence for parental effects. For all traits investigated, we did not observe a clear directional asymmetry between the reciprocal hybrid crosses or trends in the distribution of individual behavior (Fig 3, Supp. Fig. 2). Overall, only a small fraction of the variance in behaviors was attributable to differences between fathers and mothers (between 0 and 0.18, Table 2). In contrast, individual identity explained a significant part of the behavioral variation across the two rounds of measurement (adjusted R_ind_ = 0.31 to 0.38; raw R_ind_ = 0.14 to 0.43) (Supp. Fig. 1, Table 2), i.e. individual behavior is consistent (to a certain extent), despite potential effects of habituation or sensitization to handling (Fig. 3, Table 1).

**Table 2.** Repeatabilities of lab-based behaviors. Raw repeatabilities and adjusted repeatabilities after controlling for cross ID are given for individual ID, father ID and mother ID along with their 95% confidence intervals (CI)

| Behavior | Fish ID | Father ID | Mother ID |
| --- | --- | --- | --- |
|  | **R<sub>ind</sub> (95 % CI)** | **R<sub>father</sub> (95 % CI)** | **R<sub>mother</sub> (95 % CI)** |
| **Activity - Raw** | 0.14 (0.00, 0.30) | 0.08 (0, 0.25) | 0.019 (0, 0.12) |
| **Activity – Adjusted** | 0.34 (0.22, 0.53) | 0.00 | 0.18 (0, 0.40) |
| **Exploration – Raw** | 0.43 (0.29, 0.55) | 0.00 | 0.03 (0, 0.13) |
| **Exploration –Adjusted** | 0.38 (0.25, 0.54) | 0.00 | 0.10 (0, 0.27) |
| **Shoaling – Raw** | 0.31 (0.15, 0.46) | 0.03 (0, 0.13) | 0.03 (0, 0.12) |
| **Shoaling – Adjusted** | 0.31 (0.15, 0.46) | 0.00 | 0.00 |
| **Boldness – Raw** | 0.29 (0.08, 0.45) | 0.06 (0, 0.18) | 0.01 (0, 0.08) |
| **Boldness - Adjusted** | 0.33 (0.19, 0.49) | 0.05 (0, 0.19) | 0.01 (0, 0.13) |

## Discussion

We aimed to study whether genetic differentiation underlies the behavioral differentiation following habitat fragmentation in sticklebacks. Using a common garden experiment, we show that the differences between residents and migrants in shoaling and migration tendency (and to some extent also activity) have a genetic basis. In contrast, there were no clear patterns regarding differences in other behaviors or size between crosses. The earlier observed differences in these traits between wild-caught residents and migrants might therefore reflect differences in the respective developmental environments of the two ecotypes of fish. We discuss below the likely causes of divergence in our system and compare the patterns to those observed in post-glacial divergence of marine and freshwater sticklebacks. Then we discuss the eco-evolutionary implications of our findings in link with conservation plans of our study area.

Our common garden experiment revealed that the divergence in at least two of the five behavioral traits studied have a genetic basis. This corroborates a previous study on sticklebacks showing that the expression of heritable variation, i.e. the fraction of phenotypic variance owing to additive effects of genes (Lynch and Walsh 1998), substantially varied depending on the personality trait considered and the evolutionary history of the populations (Dingemanse et al., 2009). An interesting future avenue will be to quantify population specific trait heritabilities and the relative contribution of genetic and non-genetic sources of variation in those behaviors. Furthermore, it remains to be tested if the genetic differences we uncovered reflect local adaptation as opposed to other processes such as genetic drift or founder effects. Shoaling and migration tendencies are very crucial for the ancestral migratory fish. Their migratory lifestyle involves group schooling tendencies and potentially higher shoaling tendencies due to increased predator pressure owing to ‘openness’ of habitats in the sea. In residents, shoaling tendencies may be less strongly selected for, leading to the pattern of random association with the shoal that we have recovered in our experiments (Fig. 3c). Alternatively, lower shoaling tendencies may be selected for due to increased competition, for instance in winter, when resources are scarce leading to a trade-off between intra-specific aggression and competition (Lacasse and Aubin-Horth 2014). Studies on marine-freshwater stickleback pairs have also revealed potential genetic underpinnings of shoaling via *eda* gene (freshwater sticklebacks shoaled less and schooled less efficiently than migrants; Archambeault et al. 2020, Di-Poi et al. 2014, Wark et al. 2011) and migratory tendencies via genetic divergence in Thyroxine response mechanisms (Kitano et al. 2010). One next step will be to test whether the genetic differentiation of shoaling and migratory tendencies reflect local adaptation using either a genomic approach to detect signature of adaptive divergence (using, for example, a whole genome and/or a candidate gene (*eda* allele) approach) or a transplant experiment where we would raise crosses in different environmental conditions (marine vs freshwater) to infer fitness.

We expected similar differentiation in other traits, as they were found to be different between wild-caught migrants and residents over two study years (Ramesh et al. 2021). For instance, studies have shown moderately heritable and additive genetic components in behaviors such as exploration and boldness, in sticklebacks (Dingemanse et al. 2009). However, in our experiment, body size and behaviors such as exploration and boldness did not show differences between crosses. For body size, responses may be potentially plastically adjusted to the ecological conditions as seen in previous studies (for example, predation pressure; Frommen et al. 2011; niche specialization, Day and McPhail 1996; Wund et al. 2008). Similar to body size, behaviors such as exploration and boldness may also be environmentally determined. Alternatively, these behaviors could also be state-dependent (state, being size or mass in this case), owing to differences in resource availability during growth of migrants and residents (Luttbeg and Sih 2010; Wolf and Weissing 2010). It also remains possible that the differences in behavior in wild migrants are due to plastic responses of migrants in freshwater vs sea conditions, which has not been tested here.

In our current study, we found little evidence for maternal effects as maternal contribution to trait variation was small and not significant (19% for activity, 3% for exploration, and 3% for shoaling tendencies) and we did not find clear systematic differences between the reciprocal hybrid crosses (RM and MR). However, we raised the juveniles in the absence of paternal care. Hence, it remains possible that the behavioral differences observed between wild-caught migrants and residents (Ramesh et al. 2021) are related to differences in paternal care. This is an interesting avenue warranting further investigation because there is evidence for parental programming through maternal effects and paternal care in sticklebacks (Giesing et al. 2011; McGhee et al. 2012, 2015; McGhee and Bell 2014; Mommer and Bell 2014; Stein and Bell 2014).

Our studies revealing genetic differentiation between ancestral migrant and resident populations in behaviors related to migration and shoaling are timely and have important consequences for conservation efforts. Water authorities are currently implementing conservation measures which aim at restoring river connectivity via barrier removal or the construction of fishways. Reconnecting migratory and genetically differentiated land-locked populations can be viewed as a large scale eco-evolutionary experiment that raises exciting questions such as: will migratory and resident sticklebacks intermix and introgress in sympatry (Ravinet 2021)? Will hybrids be selected against, or will we have incomplete gene flow and partial migration occurring in these populations (Berner et al. 2011; Ingram et al. 2015; Hanson et al. 2016; Lackey and Boughman 2017)? From our studies, residents and hybrids show lowered migratory and shoaling tendencies. This could potentially drive divergent selection, and lead to the genetic differentiation of sympatric populations with partial migration upon reconnection. Divergence may also be maintained or enhanced by size-assortative mating of migrants and residents as size difference at maturity has been detected in the wild (Ramesh et al. 2021) or by phenotype-dependent microhabitat choice (Maciejewski et al. 2020; Dean et al. 2021). Irrespective of the mechanisms involved in the observed phenotypic differentiation between migrants and residents, whether the migrant-resident ecotype divergence will persist in the absence of migration barriers needs to be investigated.

Overall, using a common garden experiment, we found evidence for genetic differentiation in shoaling, migratory tendencies and potentially activity. These results suggest that residents may have locally adapted to their novel environmental conditions in our system. Few imminent questions that follow this finding are whether our results can be generalized to other freshwater and migratory fish species that have undergone isolation and how conservation plans may be affected (Franssen et al. 2013; Tuomainen and Candolin 2011). Conservation methods indeed should not only aim at restoring the ecosystem to its original state because this may lead to unwanted consequences (Stockwell et al. 2003). For example, reversal of responses to restorations may not be possible if newly adapted populations or species lack genetic variation, leading to a rapid population decline after conservation measures are in place (Lahti et al. 2009; Mable 2019). Alternatively, newly adapted populations or species may, in fact, have selected for invasive phenotypes such as novel foraging tactics and increased aggression and boldness, leading to unwanted expansions causing unpredictable effects on other species and communities (Holway and Suarez 1999; Sol et al. 2002). Hence conservation efforts should be aimed at implementing methods taking an informed approach of the current state of the system and assessing the evolutionary changes undergone in the species assemblages they are aimed at.

## Acknowledgments

We thank Dennis de Worst and Willem Diederik for help with fish care and advice on experimental design. We thank Peter Paul Schollema, at the Water Authorities Hunze en Aa’s and Jeroen Huisman at van Hall Larenstein, University of Applied Sciences for help with acquiring samples of sticklebacks. We thank Jakob Gismann, Jana Riederer, Raphaël Scherrer, Kevin Kort and Albertas Janulevicius for their useful comments on the manuscript. We also thank the two reviewers, who provided excellent comments and suggestions that improved the manuscript.

## Ethics approval

Sampling of wild animals and handling methods were done following a fishing permit from Rijksdienst voor Ondernemend Nederland (The Netherlands) and an angling permit from the Hengelsportfederatie Groningen-Drenthe. Animals housing and behavioral tests adhered to the project permit from the Centrale Commissie Dierproeven (The Netherlands) under the license number AVD1050020174084.

## Data availability

The final processed data used for the figures and analyses of this study are made available as supplementary material.

## Consent for publication

All authors have given their consent for publication

## Supplementary material

**Supplementary fig. 1.**
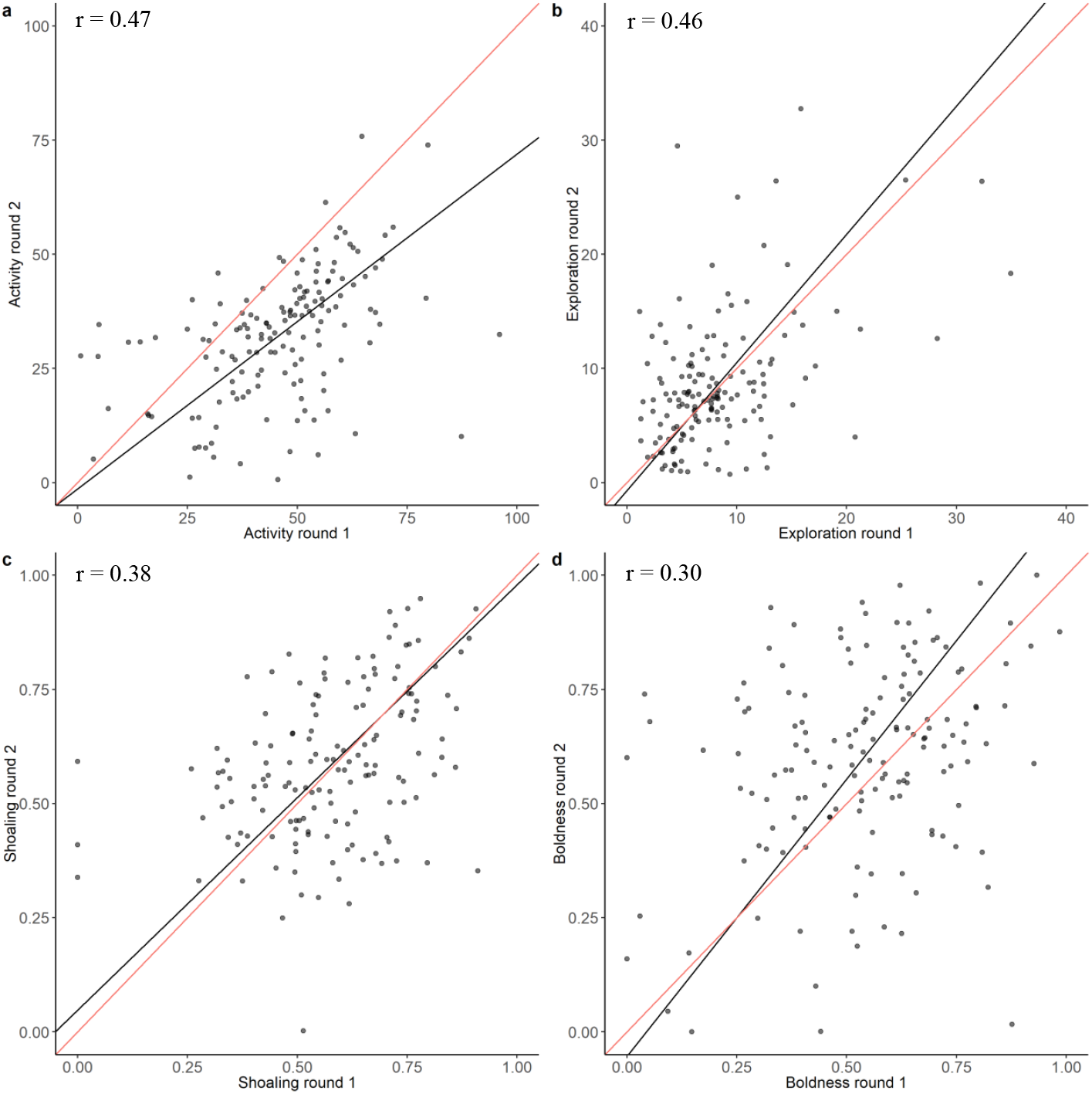
Repeatability of lab-based behaviors. Major axis regression between behaviors measured in the first and the second round. The black line is the major-axis regression line, and the red line is the main diagonal (where *y* = *x*). Pearson’s correlation coefficients are show at top left of each plot. (Sample size: N_Total_ = 151; N_MM_ = 39, N_MR_ = 38, N_RM_ = 34, N_RR_ = 40)

**Supplementary fig. 2.**
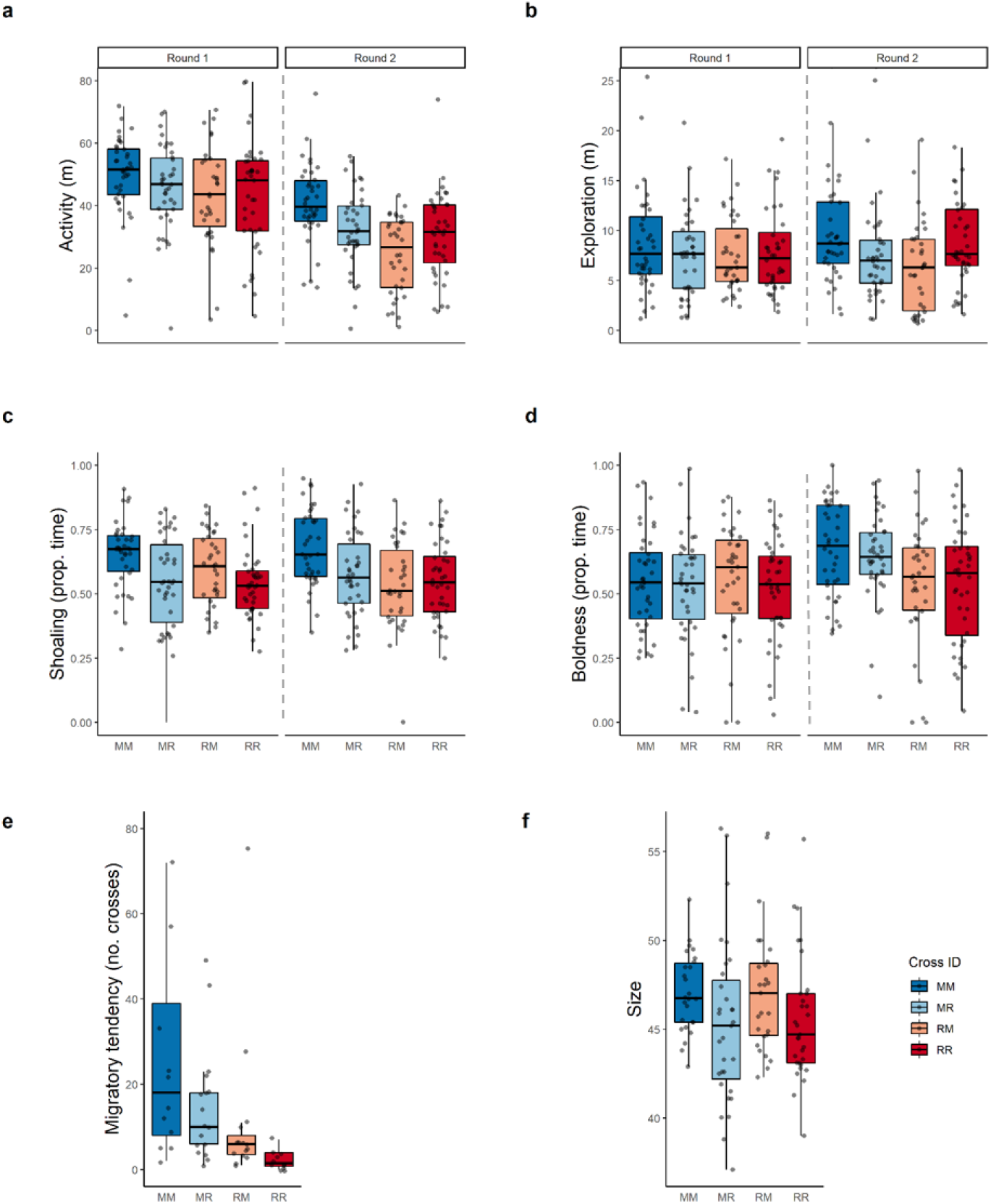
Median scores and quartiles for behaviors and size of F1 fish of different crosses along with variation within crosses. a “Activity” – total distance travelled in meters (m); b “Exploration” – total distance travelled in a novel arena in m; c “Shoaling” – Proportion of time spent near shoal compartment; d “Boldness” – Proportion of time spent near predator. For lab-based behaviors, the mean behavioral scores for the two repeats are represented separately. (Sample sizes round 1: N_MM_ = 40, N_MR_ = 39, N_RM_ = 35, N_RR_ = 40; round 2: N_MM_ = 39, N_MR_ = 38, N_RM_ = 34, N_RR_ = 40); e “Migratory tendencies” – total number of pond crosses (N_MM_ = 12, N_MR_ = 17, N_RM_ = 15, N_RR_ = 12); f “Size” – Standard length in mm (N_MM_ = 40, N_MR_ = 39, N_RM_ = 35, N_RR_ = 40)

**Supplementary table 1.** Table of pairwise significance with adjusted p-values resulting from Tukey’s HSD test using ‘emmeans’ package. Significant pair-wise comparisons are represented in bold.

| Contrast | Estimate | Std.<br>error | p value |  | Estimate | Std.<br>error | p value |
| --- | --- | --- | --- | --- | --- | --- | --- |
| Activity<br>Round 1 |  |  |  | Activity<br>Round 2 |  |  |  |
| MM-MR | 4.303 | 6.00 | 0.8867 |  | 8.965 | 6.02 | 0.4969 |
| MM-RM | 7.327 | 3.72 | 0.2526 |  | **18.531** | **3.78** | **0.0015** |
| MM-RR | 4.978 | 5.91 | 0.8329 |  | 10.039 | 5.93 | 0.4029 |
| MR-RM | 3.023 | 6.12 | 0.9577 |  | 9.566 | 6.13 | 0.4598 |
| MR-RR | 0.675 | 3.31 | 0.9968 |  | 1.074 | 3.31 | 0.9874 |
| RM-RR | -2.348 | 6.12 | 0.9791 |  | -8.491 | 6.13 | 0.5483 |
| Exploration<br>Round 1 |  |  |  | Exploration<br>Round 2 |  |  |  |
| MM-MR | 1.541 | 2.08 | 0.8774 |  | 2.741 | 2.09 | 0.5851 |
| MM-RM | 1.774 | 1.57 | 0.6798 |  | 3.977 | 1.60 | 0.1134 |
| MM-RR | 0.256 | 2.07 | 0.9993 |  | 0.573 | 2.09 | 0.9921 |
| MR-RM | 0.233 | 2.17 | 0.9995 |  | 1.236 | 2.17 | 0.9385 |
| MR-RR | -1.286 | 1.41 | 0.8003 |  | -2.168 | 1.41 | 0.4585 |
| RM-RR | -1.518 | 2.13 | 0.8894 |  | -3.404 | 2.14 | 0.4386 |
| Shoaling<br>Round 1 |  |  |  | Shoaling<br>Round 2 |  |  |  |
| MM-MR | 0.10150 | 0.0413 | 0.0979 |  | 0.09749 | 0.0416 | 0.1202 |
| MM-RM | 0.08918 | 0.0461 | 0.2383 |  | **0.16751** | **0.0474** | **0.0070** |
| MM-RR | 0.09593 | 0.0409 | 0.1135 |  | **0.11702** | **0.0408** | **0.0382** |
| MR-RM | -0.01232 | 0.0463 | 0.9932 |  | 0.07001 | 0.0470 | 0.4604 |
| MR-RR | -0.00557 | 0.0403 | 0.9990 |  | 0.01952 | 0.0402 | 0.9613 |
| RM-RR | 0.00675 | 0.0494 | 0.9991 |  | -0.05049 | 0.0496 | 0.7406 |
| Boldness<br>Round 1 |  |  |  | Boldness<br>Round 2 |  |  |  |
| MM-MR | 0.02566 | 0.0538 | 0.9627 |  | 0.02347 | 0.0545 | 0.9720 |
| MM-RM | 0.03091 | 0.0671 | 0.9660 |  | 0.11480 | 0.0680 | 0.3764 |
| MM-RR | 0.02042 | 0.0642 | 0.9882 |  | 0.13654 | 0.0648 | 0.2151 |
| MR-RM | 0.00525 | 0.0679 | 0.9998 |  | 0.09133 | 0.0681 | 0.5591 |
| MR-RR | -0.00523 | 0.0627 | 0.9998 |  | 0.11307 | 0.0627 | 0.3368 |
| RM-RR | -0.01048 | 0.0563 | 0.9976 |  | 0.02174 | 0.0566 | 0.9797 |

